# COVID-19 mRNA vaccines drive differential Fc-functional profiles in pregnant, lactating, and non-pregnant women

**DOI:** 10.1101/2021.04.04.438404

**Authors:** Caroline Atyeo, Elizabeth A. DeRiso, Christine Davis, Evan A. Bordt, Rose M. DeGuzman, Lydia L. Shook, Lael M. Yonker, Alessio Fasano, Babatunde Akinwunmi, Douglas A. Lauffenburger, Michal A. Elovitz, Kathryn J. Gray, Andrea G. Edlow, Galit Alter

## Abstract

Significant immunological changes occur throughout pregnancy to tolerize the mother and allow growth of the fetal graft. However, additional local and systemic immunological adaptations also occur, allowing the maternal immune system to continue to protect the dyad against foreign invaders both during pregnancy and after birth through lactation. This fine balance of tolerance and immunity, along with physiological and hormonal changes, contribute to increased susceptibility to particular infections in pregnancy, including more severe COVID-19 disease. Whether these changes also make pregnant women less responsive to vaccination or induce altered immune responses to vaccination remains incompletely understood. To holistically define potential changes in vaccine response during pregnancy and lactation, we deeply profiled the humoral vaccine response in a group of pregnant and lactating women and non-pregnant age-matched controls. Vaccine-specific titers were comparable, albeit slightly lower, between pregnant and lactating women, compared to non-pregnant controls. Among pregnant women, we found higher antibody titers and functions in those vaccinated with the Moderna vaccine. FcR-binding and antibody effector functions were induced with delayed kinetics in both pregnant and lactating women compared to non-pregnant women. Antibody boosting resulted in high FcR-binding titers in breastmilk. These data point to an immune resistance to generate highly inflammatory antibodies during pregnancy and lactation, and a critical need to follow prime/boost timelines in this vulnerable population to ensure full immunity is attained.

## Introduction

Pregnant women experience both increased disease severity and morbidity upon SARS-CoV-2 infection *(1, 2)*. However, pregnant and lactating women were left out of initial COVID-19 vaccine trials due to heightened safety concerns *(3–6)*. Given that pregnant women are vulnerable to severe SARS-CoV-2 disease, it is important to understand the immunological response to vaccination in pregnant and lactating women. Understanding how pregnancy/lactation affects response to vaccination and antibody transfer to infants offers critical opportunities to guide recommendations for this population.

Pregnant and lactating women are routinely encouraged to receive vaccines against influenza and pertussis *(7, 8)*. Mounting data point to dampened vaccine-induced antibody responses in pregnant women marked by a lower fold increase in antibody titers, lower neutralizing antibody responses, and reduced T cell immune responses compared to non-pregnant women *(9)*. Given the significant morbidity associated with influenza infection in pregnant women annually, influenza vaccination is recommended throughout gestation *(10)*, whereas pertussis vaccination is recommended in the late second and early third trimester, to facilitate maximal antibody transfer and protect the developing neonate *(11–13)*. However, because most women have been exposed or previously immunized against these pathogens, these vaccines largely boost immunity, rather than prime a de novo immune response. Thus, whether the same principles for antibody transfer will apply to novel vaccine platforms used against SARS-CoV-2, as well as to a new antigen (SARS-CoV-2 spike), remains unclear. Moreover, whether vaccine-induced immune profiles will vary across pregnancy and lactation, impacting antibody transit across the placenta or into breastmilk, is not known. This understanding could provide crucial insights to guide the optimal administration of the vaccine to women and their infants.

The first two vaccines that were approved for emergency use authorization (EUA) by the Food and Drug Administration (FDA) are novel vaccine platforms that use mRNA to induce an immune response against the SARS-CoV-2 spike. Linked to highly effective protection against severe COVID-19 disease in non-pregnant populations *(14, 15)*, both mRNA platforms clearly lead to the induction of robust immunity in men and non-pregnant women across age groups (Anderson *et al.*, 2020; Walsh *et al.*, 2020). Moreover, emerging data suggest that mRNA vaccines also induce comparable antibody titers and neutralization in pregnant and lactating women, linked to transfer of antibodies to neonates *(16, 17)*. The explosion of multiple vaccine platforms, coupled with the availability of novel high-dimensional antibody profiling technologies, enables the unprecedented opportunity to dissect the de novo mRNA vaccine-induced immune response in this vulnerable population.

In addition to the role of antibodies in binding and neutralization, antibodies contribute to protection against COVID-19 disease through their ability to recruit the innate immune response with their Fc-domain *(18, 19)*. The Fc-functions of the humoral immune response play a critical role in resolution of COVID-19 *(20)*, are associated with protection from infection following vaccination *(21)*, play a critical role in antibody transfer across the placenta *(22–25)* and may also influence transfer into breastmilk *(26)*. To define qualitative features of the vaccine-induced humoral immune response across pregnancy and early life, here we comprehensively profiled the humoral immune response following mRNA vaccination in pregnant, lactating, or non-pregnant women who received the Pfizer/BioNTech or the Moderna vaccine. While overall antibody titers were comparable, we observed reduced Fc-receptor (FcR)-binding and functional antibody evolution in pregnant and lactating women that required a boost to reach full functional capabilities. Compromised placental transfer was observed, likely due to vaccination close in timing to delivery, with improved transfer noted with increased time from immunization. Robust levels of IgG and IgA were noted in breastmilk. Differences were observed across vaccine platforms, marked by differential skewing of IgG and IgA responses in blood and milk. These data point to significant differences in vaccine-induced antibody profiles among pregnant, lactating and non-pregnant women that influence the level and quality of immune transfer to neonates, clearly arguing for a need for timely boosting in this vulnerable population and a a need to understand how timing of vaccine administration in pregnancy impacts maternal immune response and placental antibody transfer.

## Results

### The vaccination-induced antibody response in pregnant, lactating, and non-pregnant women

Two mRNA vaccines were the first EUA approved vaccines, showing ~95% protection against severe COVID-19 disease *(14, 27–29)*. Emerging data have begun to illustrate the robust immunogenicity of these vaccines in pregnant and lactating women, in the absence of enhanced reactogenicity *(16)*. However, whether the overall humoral immune profile diverges in pregnant or lactating women and if these profiles impact transfer to neonates remains incompletely understood. Thus, the SARS-CoV-2 humoral immune response was characterized in a cohort of 84 pregnant, 31 lactating and 16 non-pregnant age-matched controls. Individuals were sampled before vaccination (V0), after first vaccination (prime, V1) and/or after second vaccination (boost, V2).

After the prime, clear differences were noted between serum antibody responses of pregnant/lactating women and non-pregnant women (**Figure 1A**). Most differences related to lower antibody titers after the prime (PC1) and Fc-receptor (FcR) binding capacity (PC2) among pregnant and lactating women compared with non-pregnant women. After the boost, there were no significant differences between pregnant/lactating and non-pregnant women (**Figure 1B**). Although some differences persisted, differences were nearly exclusively linked to enhanced FcR-binding in non-pregnant women.

**Figure 1.**
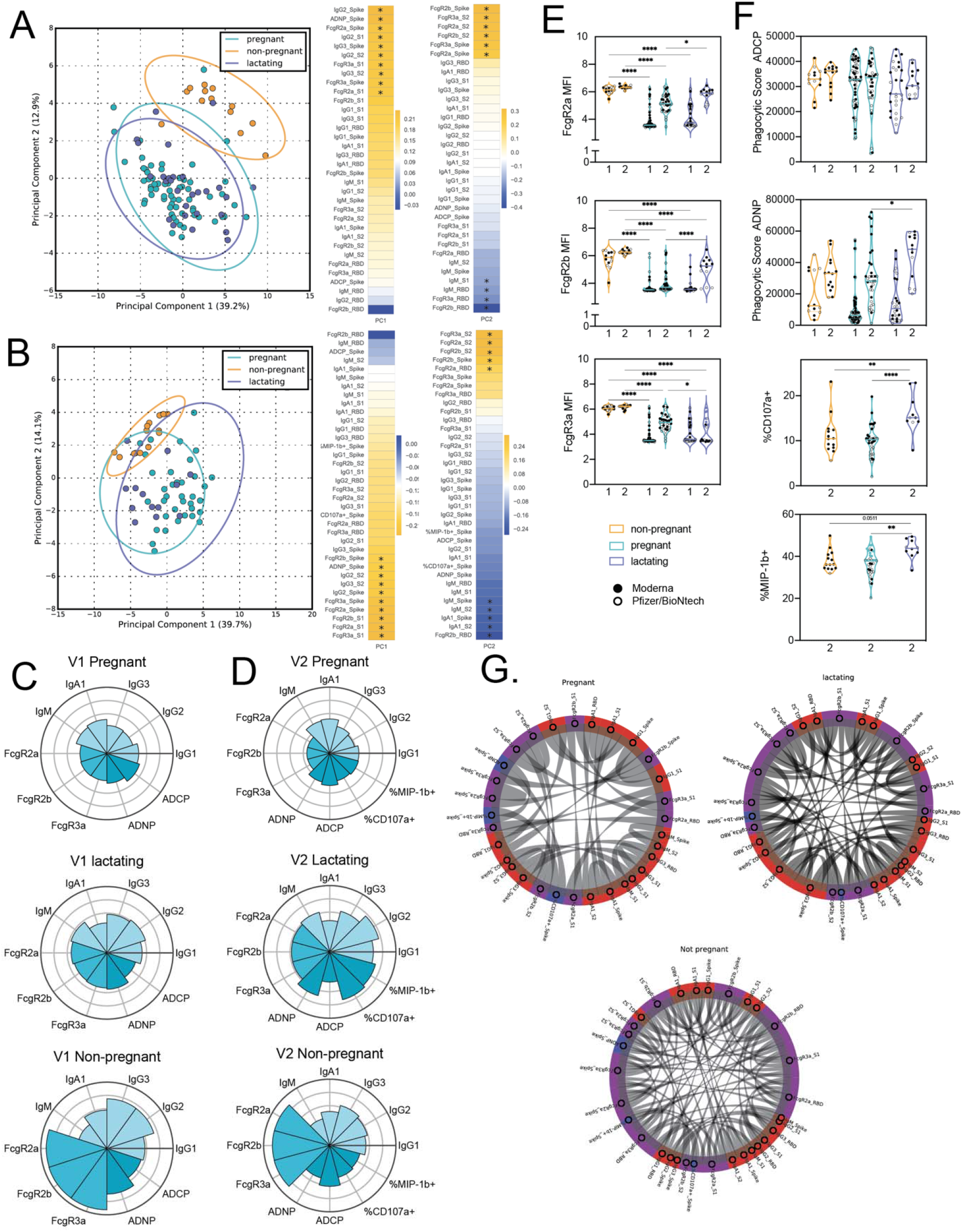
Vaccination induces enhanced FcR-binding in non-pregnant women. **A-B.** A principal component analysis (PCA) was built using measured antibody features after V1 (A) or V2 (B). The dot plots show the scores of each individual, with each dot representing an individual. The ellipses represent the 95% confidence interval for each group. The heatmaps show the features that contribute to the loadings along each principal component (PC). An asterisk (*) indicates that the corresponding feature was among the top 10 features that contributed to the loadings along the PC indicated. **C-D**. The polar plots show the mean percentile rank for each feature for V1 (C) and V2 (D) samples. Features were ranked separately for each time point. **E**.The violin plots show the FcγR-binding for non-pregnant, pregnant, and lactating women at V1 (1) and V2 (2). The filled dots show the titer for women who received the Moderna vaccine, and outlines show the titer for women who received the Pfizer/BioNtech vaccine. Significance was only calculated between groups in the same time point and was determined by a one-way ANOVA with followed by posthoc Šidák’s multiple comparison test. * p <0.05, ** p < 0.01,*** p < 0.001, **** p < 0.0001. **F**.The violin plots show the antibody functions for non-pregnant, pregnant, and lactating women at V1 (1) and V2 (2). The filled dots show the titer for women who received the Moderna vaccine, and outlines show the titer for women who received the Pfizer/BioNtech vaccine. Significance was determined by a one-way ANOVA followed by posthoc Šidák’s multiple comparison test. * p <0.05, ** p < 0.01,*** p < 0.001, **** p < 0.0001. **G**.Increased coordination in the antibody response of non-pregnant women. The chord diagrams connect the features that have a spearman correlation > 0.75 and Bonferroni-corrected p-value > 0.05 for non-pregnant, pregnant, and lactating women.

To further understand the difference in individual antibody features in pregnant, lactating, and non-pregnant women, we plotted the mean percentile rank of each spike (S)-specific feature measured at V1 and V2 (**Figure 1C-D**). After the prime (V1), non-pregnant women had higher IgG subclass responses, higher antibody functions and higher FcR-binding compared to pregnant and lactating women (**Figure 1C**). At this time point, pregnant and lactating women had a similar antibody response. Interestingly, after the boost (V2), lactating women boosted their antibody response more effectively than pregnant women, marked by higher IgG-levels and higher NK-cell activity (%CD107a+ and %MIP-1b+) (**Figure 1D**). After boost, the vaccine response in lactating women was similar to that of non-pregnant women, although lactating women had lower FcR-binding compared to non-pregnant women, who maintained higher FcR-binding levels after the boost.

Next, to specifically capture the differences in the quality of the vaccine induced humoral immune response across the women, FcR-binding and Fc-effector profiles were compared at a univariate level across the groups. Strikingly, FcR-binding antibodies across all FcRs were significantly higher in non-pregnant compared to pregnant and lactating women at V1 (**Figure 1E**). However, the boost raised these antibody responses in both pregnant and lactating women, albeit to lower levels than observed in non-pregnant women (**Figure 1E**). Slight, but largely insignificant, differences were noted in antibody isotype and subclass evolution across the groups (**Supplemental Figure 1**), with small rises in antibody levels following boosting. Interestingly, whereas antibody-dependent cellular phagocytosis (ADCP) was induced at similar levels in the three populations and was not increased by the vaccine boost, antibody-dependent neutrophil phagocytosis (ADNP) was similar between groups after the prime and was augmented in all 3 groups after boosting, with the most significant boost in lactating women (**Figure 1F**). The ability of antibodies to drive NK cell activation (CD107a, a degranulation marker and MIP-1β, a chemokine) was distinct across the groups, whereby lactating women induced significantly higher NK cell activating antibodies after boosting compared to pregnant and non-pregnant women (**Figure 1F**). Overall, these higher FcR-binding profiles in non-pregnant and lactating women were linked to enhanced coordination in the humoral immune responses compared to pregnant women, the latter showing sparser coordination in the vaccine induced humoral immune response (**Figure 1G**). These data reveal that pregnant and lactating women show potential early deficiencies in their vaccine induced immune response, that recover at boosting in lactating women and to a lesser extent in pregnant women.

### Differences in maternal serum vs cord blood antibodies

Previous studies focused on pertussis vaccination have pointed to the selective and active transfer of highly functional antibodies across the placenta, marked by the specific selective transfer of FcγR3a binding antibodies *(23)*. However, more recent studies of SARS-CoV-2 infection in pregnancy have noted compromised transfer with infection in the third trimester, linked to reduced antibody transfer (transfer ratio < 1) but maintaining a selection bias based on binding to FcγR3a *(25)*. To begin to understand the overall profiles of antibodies that are transferred from pregnant individuals to infants, we profiled the humoral immune responses across the maternal and umbilical cord blood. Overall, higher levels of antibodies were observed in maternal blood compared to cord blood (**Figure 2A**). Variable patterns of transfer of IgG titer, FcR-binding and antibody function were observed from the mother to the cord (**Figure 2B-C**). Despite the recency of vaccination, equivalent, IgG1 spike (S)-specific titers were transferred across the placenta to the infant (**Figure 2A**). Lower IgG3 transfer was noted. Lower FcR-binding antibodies were transferred. Despite previous observations of augmented NK cell activating antibody transfer following vaccines that boost previously established immunity, such as to pertussis and influenza *(23)*, stable phagocytic antibodies but decreased NK-cell activating antibody were transferred to infants (**Figure 2B**). Conversely, active transfer of more functional NK cell activating antibodies was noted for influenza hemagglutinin (HA)-specific antibodies in the same mother:cord pairs (**Supplemental Figure 2A-B**) suggesting that reduced transfer of S-specific antibodies is not attributable to vaccine-induced changes in placental activity. Instead, decreased S-specific transfer could be linked to time from vaccination (**Supplemental Figure 2C-D**), suggesting that vaccination proximal to the time of birth may simply not permit the effective transfer of the most functional antibody subpopulations.

**Figure 2.**
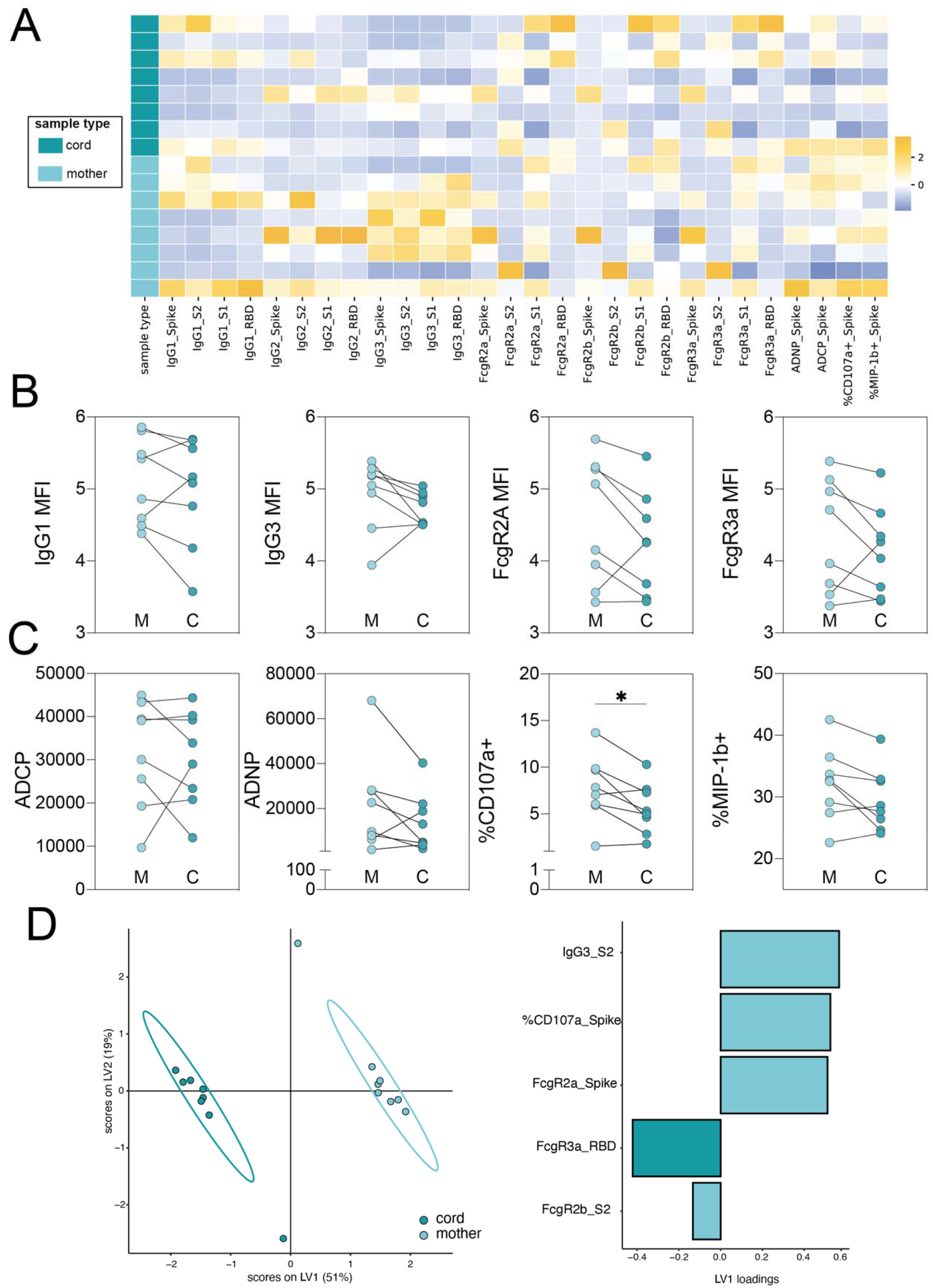
Vaccination results in the transfer of FcR-binding antibodies into the cord blood. **A.** The heatmap shows the z-scored SARS-CoV-2-specific data for each cord and matched mother. Each row represents an individual sample. **B.** The dot plots show the IgG1, IgG3, FcγR2a-binding and FcγR3a-binding titer against spike for maternal (M) and cord (C) blood. Lines connect maternal:cord dyads. Significance was determined by Wilcoxon-matched pairs signed rank test. * p <0.05, **p < 0.01, ***p < 0.001, **** p < 0.0001. **C.** The dot plots show the ADCP, ADNP and ADNKA (CD107a and MIP-1β) functional titer against spike for maternal (M) and cord (C) blood. Lines connect maternal:cord dyads. * p <0.05, **p < 0.01, ***p < 0.001, **** p < 0.0001. **D.** A multilevel partial least-squares discriminant analysis (mPLSDA) was built using maternal or cord LASSO-selected SARS-CoV-2-specific features. The dot plot (left) shows the scores of each sample, with the ellipses representing the 95% confidence interval for each group. The bar plot (right) shows the loadings of each LASSO-selected feature for the mPLSDA. The color of the bar indicates in which group the feature is enriched.

Given the de novo nature of the response to SARS-CoV-2 S and the novelty of this vaccine platform, we next aimed to determine whether placenta transfer was strictly governed by total amounts of antibody or based on specific characteristics of the vaccine-induced humoral immune response. Thus, we performed a multilevel partial least squares discriminant analysis (mPLSDA) to determine the S-specific antibody features that transferred preferentially across the cord (**Figure 2D**, right). Fc-profiles in maternal and cord blood were completely distinct, with expected higher overall levels of antibodies in maternal blood, as the vaccine was only administered in these dyads in the third trimester. However, despite the lower levels of antibodies in the cord, receptor binding domain-specific FcγR3a binding was enriched in the cord (**Figure 2D**, left). Thus, similar to the previously observed transfer sieve, even at low antibody-transfer rates, the placenta selects for FcγRIIIa-binding, functionally enhanced vaccine-induced antibodies aimed at enriching infants with the most protective antibodies.

### Optimal vaccine-induced antibody transfer to the breastmilk requires boosting

Beyond placental transfer, antibody transfer can continue to occur after birth through breastmilk. Vaccine immunity changes over time in immunized lactating women, but how this influences antibody transfer to the infant is incompletely understood. Using an unsupervised PCA of the post-prime and post-boost vaccine induced immune response, we observed a slightly expanded antibody functional and FcR-binding response in the serum of lactating women after the boost (**Figure 3A**), pointing to a functional maturation of the humoral immune response with boosting. Similarly, in breastmilk, boosting resulted in the robust transfer of FcR-binding antibodies and highly functional IgG3 antibodies (**Figure 3B and Supplemental Figure 3A**). To gain specific insights into the antibody subpopulations that are transferred most efficiently across the blood and breastmilk, we next plotted the mean percentile rank of each spike-specific antibody feature for the prime and boost (**Figure 3C-D**). The polar plots highlight the preferential boosting of FcR-binding IgG responses in the serum following boosting, with a prominent expansion of IgG and FcR binding, but only a marginal impact on enhancing IgA and IgM responses (**Figure 3C**). Moreover, the same transfer profile was noted in the breastmilk (**Figure 3D**), with a high transfer of IgGs with FcR-binding capabilities after the boost. To further understand the sieve of antibodies from serum to breastmilk upon vaccination, we plotted the transfer ratio (breastmilk MFI/serum MFI) of isotypes and FcRs against the SARS-CoV-2 spike after the first dose (**Figure 3E**) and after the boost (**Figure 3F and Supplemental Figure 3B**). After the prime and the boost, IgA was the most preferentially transferred of any isotype. Interestingly, IgG2 was transferred more highly to breastmilk than any other IgG subclass after the first dose, but after the boost, IgG3 was the most highly transferred subclass. Moreover, after the first dose, there was a preferential transfer of antibodies that could bind FcγR3a, whereas antibodies that could bind FcγR2b, the inhibitory FcR, had the lowest transfer ratio. After the boost, however, there were no significant differences in the transfer of FcR-binding antibodies, pointing to robust transfer of all FcR-binding antibodies to breastmilk after a boost. In addition, we analyzed the transfer ratio of antibody functions after prime and after boost. Whereas ADCP and ADNP were transferred at equivalent ratios after prime (**Figure 3G**), antibodies able to drive ADCP were transferred at a higher ratio than those able to activate ADNP (**Figure 3G**), likely reflecting the enhanced ADNP activity in the serum of lactating women after boost. In addition, NK-cell activating antibodies were transferred at low levels post-boost (**Supplemental Figure 3C**), suggesting a sieve at the mammary gland, preventing the transfer of highly inflammatory antibodies through breastmilk. This analysis revealed enhanced functional antibodies in breastmilk following the boost, accompanied by decreased IgM and IgA induction post-boost. Collectively, these data emphasize that while the breast clearly enriches IgM and IgA delivery to the breastmilk *(30, 31)*,vaccination appears to dramatically augment highly functional IgG transit to the milk that are likely key to antiviral immunity across viral pathogens *(31)*.

**Figure 3.**
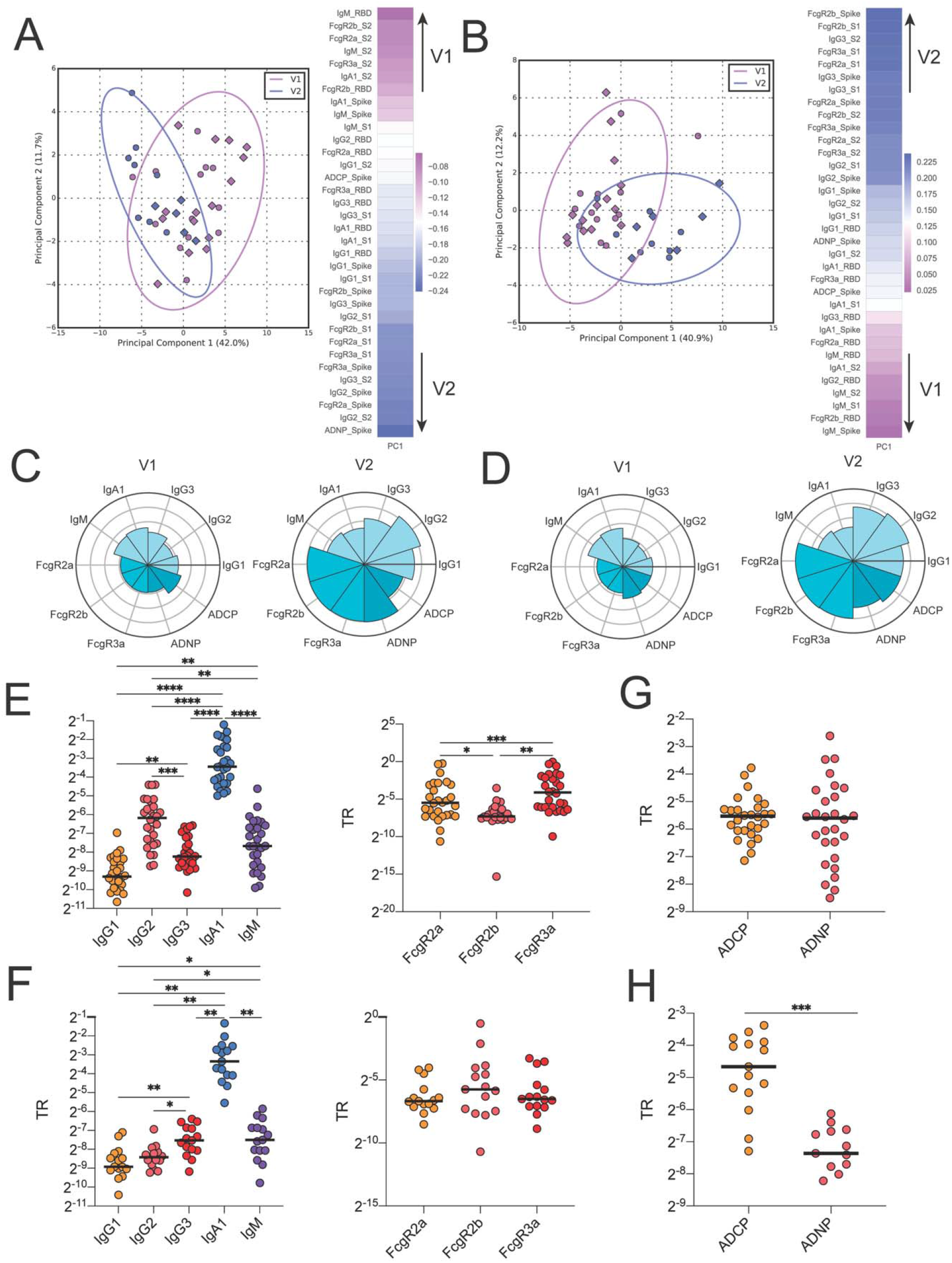
Second dose of vaccination required for boost in functional antibody response in serum and breastmilk of lactating women. **A**. A PCA was built on the SARS-CoV-2-specific features at V1 (purple) and V2 (blue) in lactating women. The ellipses represent the 95% confidence interval for each group. The heatmaps show the contribution of each feature along each principal component (PC). The color of the heatmap indicates in which group each feature is enriched. **B**. A PCA was built on the SARS-CoV-2-specific features in breastmilk at V1 (purple) and V2 (blue), using time point as the outcome variable. The ellipses represent the 95% confidence interval for each timepoint. The heatmap shows the contribution of each feature to the corresponding PC. **C**. The polar plots show the mean percentile rank for each feature for V1 or V2 in the serum of lactating women. The color of the heatmap indicates in which group each feature is enriched. D. The polar plots show the mean percentile rank for each feature for V1 or V2 in breastmilk. E. The dot plot shows the transfer ratio (breastmilk/serum) for each isotype (left) and FcR (right) measured at V1. Significance was determined by a one-way ANOVA followed by Tukey’s multiple comparison correction. * p <0.05, ** p < 0.01, *** p < 0.001, **** p < 0.0001. F. The dot plot shows the transfer ratio (breastmilk/serum) for each isotype (left) and FcR (right) measured at V2. Significance was determined by a one-way ANOVA followed by Tukey’s multiple comparison correction. * p <0.05, ** p < 0.01,*** p < 0.001, **** p < 0.0001. G. The dot plots show the transfer ratio (TR) (breastmilk/serum) between ADCP and ADNP measured at V1. Significance was determined by Wilcoxon-matched pairs signed rank test. * p <0.05, ** p < 0.01,*** p < 0.001, **** p < 0.0001. H. The dot plots show the transfer ratio (TR) (breastmilk/serum) between ADCP and ADNP measured at V2. Significance was determined by Wilcoxon-matched pairs signed rank test. * p <0.05, ** p < 0.01,*** p < 0.001, **** p < 0.0001.

### Moderna and Pfizer vaccination induce differential antibody responses

While both the Moderna and Pfizer vaccines exploit mRNA-based technologies, differences in mRNA dosage, lipid carriers, and vaccine dosing regimens may alter the quality of the humoral immune response across the vaccines. Thus, we compared immune responses across all women dosed with these mRNA vaccines. Although minimal differences were noted across the vaccine-induced immune responses post-prime (V1) (**Supplemental Figure 3A**), separation was observable in the vaccine-induced antibody profiles following the boost (V2) (**Figure 4A**). Notably, Moderna mRNA vaccinated women exhibited enriched neutrophil activating antibodies (ADNP), higher levels of vaccine-specific IgA, IgG2, IgG3, and NK cell activating antibodies (CD107a) compared to Pfizer vaccinated women (**Figure 4A, Supplemental Figure 3B**). Conversely, women receiving the Pfizer vaccine exhibited a slight enrichment of IgG1-and FcγR3A (**Figure 4A**) binding humoral immune responses.

**Figure 4.**
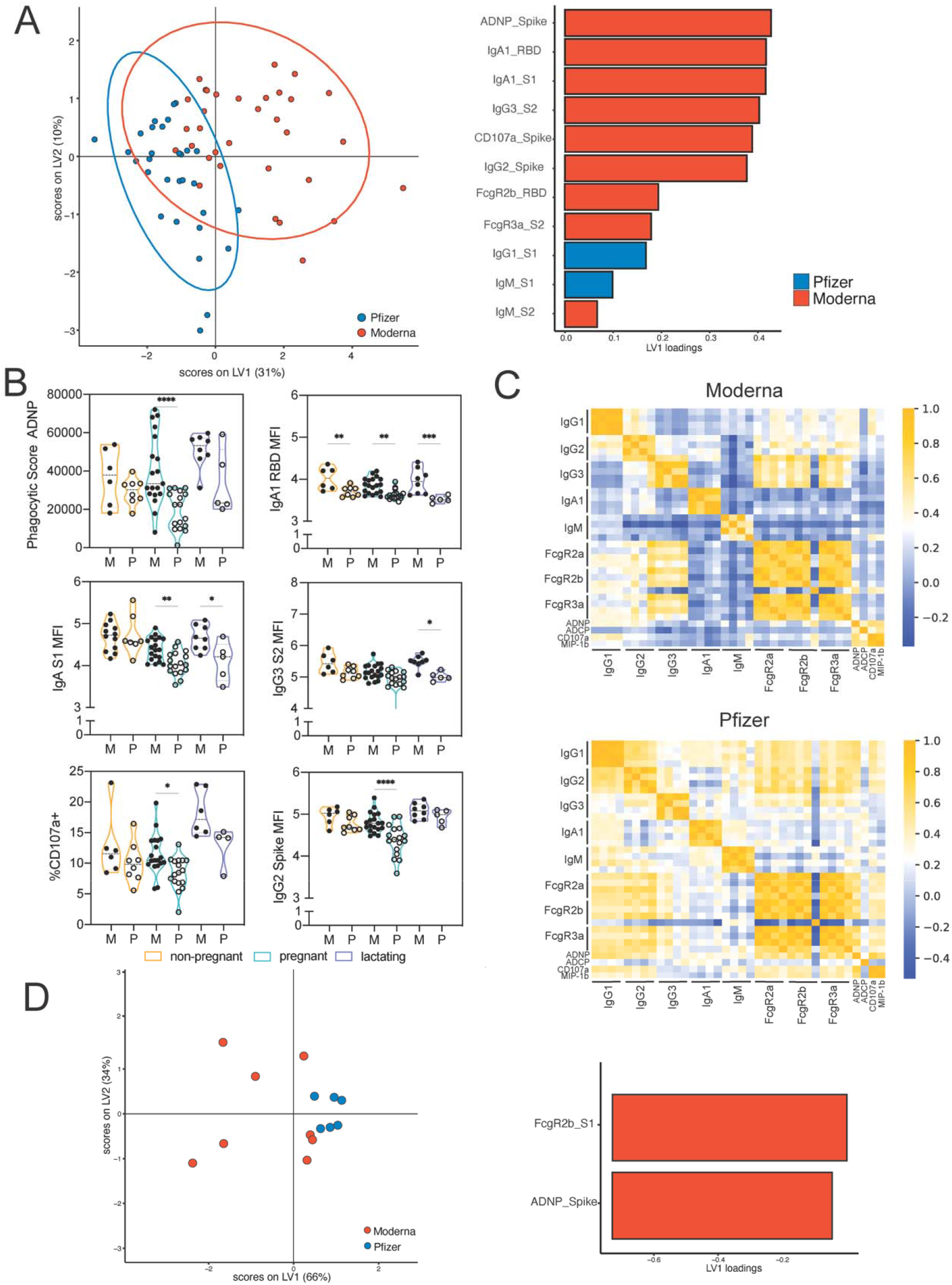
Differences in the antibody response to Moderna and Pfizer vaccination. A. LASSO-PLSDA model was built on V2 data from all groups. The dot plot (top) shows the scores for each sample, with each dot representing a sample. The ellipses represent the 95% confidence interval for each group. The bar plot (bottom) shows the loadings of each LASSO-selected feature, where the color marks the group enrichment. B. The violin plots show differences in the top LASSO-selected features from Figure 4A in non-pregnant, pregnant and lactating women given either the Moderna (M, filled dots) or Pfizer/ BioNtech (P, outline dots). All data in the violin plots is from V2. Significance was only calculated between groups in the same time point and was determined by a one-way ANOVA with followed by posthoc Šidák’s multiple comparison test. * p <0.05, ** p < 0.01,*** p < 0.001, **** p < 0.0001. C. The heatmaps show the spearman correlation between features for women who received the Moderna (top) or Pfizer (bottom) vaccine. For isotypes and FcγR-binding, the features are ordered as Spike, S2, S1, and RBD. For functional data, only functional activity against Spike was measured. D. A LASSO-PLSDA model was built on V2 breastmilk data. The dot plot (top) shows the scores for each sample, with each dot representing a sample. The ellipses represent the 95% confidence interval for each group. The bar plot (bottom) shows the loadings of each LASSO-selected feature, where the color marks the group enrichment.

To further dissect these differences, women were split into groups by their pregnancy and lactation status. While differences in IgA levels were noted across all groups, differences in ADNP, and NK cell recruiting antibodies were amplified in pregnant and lactating women (**Figure 4B**), with a dramatic expansion of functional antibodies in women following the Moderna mRNA vaccine. Moreover, to further understand the functional basis for these differences in function, we examined the functional coordination of the humoral immune response induced by each vaccine across the 3 populations of women (**Figure 4C**). Strikingly, Moderna vaccination resulted in a more focused functional coordination in the humoral immune response, centered around a highly functional IgG1/IgG3 response with robust FcR-binding and functional coordination. Conversely, women receiving the Pfizer vaccine generated a broader coordinated immune response including IgG2 and IgM responses and the exclusion of monocyte phagocytosis (ADCP), potentially pointing to a more diffuse overall humoral immune coordination profile. These serum differences translated to differences in breastmilk transfer (**Figure 4D**), with enhanced FcR-binding antibody and neutrophil-functional antibody transfer to breastmilk observed in Moderna-immunized lactating women. Whether these differences are attributable to dose, lipids, or timing of vaccination remains unclear, but provide the first clues that mRNA platforms may be selectively deployed to enhance protection in neonates once precise mechanistic correlates of immunity are defined.

## Discussion

Both EUA-approved COVID-19 mRNA vaccines have been shown to be safe and highly immunogenic in non-pregnant populations *(14, 15)*, and emerging data suggest that the vaccines are immunogenic and similarly reactogenic in pregnant and lactating women *(16)*. However, pregnancy and lactation represent distinct immunological states *(32, 33)*, that have been previously associated with reduced immunogenicity *(9)*. Whether this unique immune state is associated with the evolution of distinct humoral immune profiles upon vaccination remains incompletely understood. Using systems serology, we observed changes in the magnitude, kinetics, and quality of functional profiles of vaccine-induced antibodies. We additionally found differences in the overall antibody profile across women receiving the Moderna and Pfizer/BioNtech vaccines. These findings collectively point to an extended window of vulnerability in pregnancy/lactation following vaccination, requiring timely boosting to achieve fully functional matured antibodies to empower the pregnant individual and their child.

Pregnancy represents a delicate immunological balance which has been associated with enhanced vulnerabilities to infection in pregnant women who experience more severe influenza infection and SARS-CoV-2 infection *(34, 35)*. This vulnerability has been linked to dampened, rather than blocked, pro-inflammatory immunity, with reduced responsiveness to vaccination *(9)*. Beyond quantitative measures of pregnancy associated changes, less is known about the qualitative functional changes in the vaccine-induced humoral response during pregnancy. Here, we observed a delay in the evolution of FcR-binding and functional antibody responses in pregnant and lactating women after initial vaccination with a de novo pathogen. Conversely, we observed higher levels of functional antibodies for NK cell activity and neutrophil phagocytosis in lactating women compared to both pregnant and non-pregnant women upon boost. These data point to distinct response profiles across each of these immunological states, raising the possibility that vaccines may drive significantly different antibody functional profiles, programmed evolutionarily to maximize protection for the mother-baby dyad in that unique immune state. Given the low responsiveness to vaccination after prime, these data also highlight the critical importance to adhere to vaccination boosting among this population to optimize immunity in pregnant and lactating women.

Vaccination during pregnancy increases the passive protection transferred to newborns, who are at increased risk for severe disease upon infection due to their immature immune system. Although few newborns have been infected with SARS-CoV-2, those who have been infected have more severe outcomes than older children *(36–38)*. In the case of natural infection, poor placental transfer of antibodies has been observed in women infected in the third trimester, but these transfer ratios increase significantly to expected levels above a transfer ratio of 1 in women infected earlier in pregnancy *(39, 40)*. Likewise, women in our study that gave birth were all immunized in the third trimester and had transfer ratios above one, likely reflective of the proximity of vaccination to delivery. As more women vaccinated in the second trimester and earlier go on to deliver, it will be important to determine whether the COVID vaccines, which induce novel immune responses to pathogens never seen by our immune systems, may require administration even earlier in pregnancy than vaccines that evoke recall responses to offer optimal immunity to the neonate. In addition to antibodies transferred through the placenta, antibodies transferred through breastmilk have been shown to play a role in the protection against respiratory infections during early life *(41–43)*. Interestingly, breastmilk antibodies are highly dependent on the second dose of the vaccine to boost the transfer of functional, FcR-binding antibodies. Previous work has shown that IgGs in breastmilk may provide protection against viral infection in early life *(44–46)*. Defining the mechanism driving breastmilk transfer of IgGs could lay the foundation for designing next generation vaccines able to provide global protection for infants following birth.

Despite the delayed kinetics and functional antibody responses in pregnant versus non-pregnant women, pregnant and lactating women generated more functional neutrophil and NK cell recruiting antibodies after receiving the Moderna vaccine compared to the Pfizer/BioNtech vaccine. These enhanced functions were accompanied by a more restrictive, coordinated humoral immune response compared to women who received the Pfizer/BioNTech vaccine. Whether this is related to differences in the dose, the lipid carriers, or the dosing window (4 vs. 3 weeks) remains unclear. The extra week prior to boosting may provide the time needed for the humoral immune response to mature, resulting in more functional antibody profiles. Whether the functional advantage described here in the Moderna-generated humoral immune profile results in improved clinical protection against COVID-19 remains to be determined. Critically, optimal dosing and intervals may vary across populations and should be based on empirical data, strongly arguing for the importance of research in pregnancy and lactation to protect this vulnerable population who are often neglected during vaccine development.

Overall, our data have demonstrated that while antibody titers are similar, pregnant and lactating women respond to vaccination in qualitatively and kinetically distinct manners compared to non-pregnant women. While this study largely profiled responses in women vaccinated later in pregnancy, these data point to the importance of profiling women receiving COVID-19 vaccines throughout pregnancy to begin to understand how distinct platforms, vaccines, populations, and timing affect the quality and quantity of immunity induced across the mother:fetal dyad. Collectively, these data highlight to the importance of defining the immunology of pregnancy to develop evidence-based recommendations for vaccine recommendations and to help inspire the development of vaccines and therapeutics that may act more effectively for this unique population where optimal immunological responses are necessary to protect both mother and fetus.

## Materials and Methods

### Study participants

Women at two tertiary care centers were approached for enrollment in an IRB-approved (protocol #2020P003538) COVID-19 pregnancy and lactation biorepository study between December 17, 2020 and February 23, 2021. Eligible women were: (n=84 pregnant; n=31 lactating; or n=16 non-pregnant and of reproductive age (18-45); greater than or equal to 18 years old, able to provide informed consent, and receiving the COVID-19 vaccine.

### Study Eligibility

Eligible study participants were identified by practitioners at the participating hospitals or were self-referred. A study questionnaire was administered to assess pregnancy and lactation status, history of prior SARS-CoV-2 infection, timing of COVID-19 vaccine doses, type of COVID-19 vaccine received (BNT162b2 Pfizer/BioNTech (n= 65) or mRNA-1273 Moderna/NIH (n= 66)).

### Sample Collection

Blood and breastmilk from lactating women were collected at: V0 (at the time of first vaccine dose/baseline), V1 (at the time of second vaccine dose/ “prime” profile), V2 (2-5.5 weeks following the 2nd vaccine dose/ “boost” profile), and at delivery (for pregnant participants who delivered during the study timeframe). Umbilical cord blood was also collected at delivery for pregnant participants. The V2 timepoint reflects full antibody complement, achieved one week after Pfizer/BioNTech and two weeks after Moderna/NIH10,11. Blood was collected by venipuncture (or from the umbilical vein following delivery for cord blood) into serum separator and EDTA tubes. Blood was centrifuged at 1000g for 10 min at room temperature. Sera and plasma were aliquoted into cryogenic vials and stored at −80°C.

### Antigens

Antigens used for functional and Luminex based assays were: SARS-CoV-2 RBD (Sino Biological), SARS-CoV-2 Spike (LakePharma), SARS-CoV-2 N (Aalto Bio Reagents) and a mix of HA A/Michigan/45/2015 (H1N1), HA A/Singapore/ INFIMH-16-0019/2016 (H3N2), HA B/Phuket/3073/2013 (Immunetech). Antigen was biotinylated using Sulfo-NHS-LC-LC biotin (Thermo) and desalted using Zeba Columns (Thermo).

### Cell lines

THP-1 cells used in phagocytic assays were grown in RPMI media supplemented with 10% FBS, 5% penn/strep, 5% L-glutamine, 5% HEPES buffer (pH 7.2) and 0.5% 2-Mercaptoethanol, and maintained at 2.5×105 cells/ml.

### Primary Cells

Human neutrophils and NK cells were isolated from fresh peripheral blood. Peripheral blood was collected by the MGH Blood Bank or by the Ragon Institute from healthy volunteers. All volunteers were over 18 years of age and gave signed consent. Samples were deidentified before use. The study was approved by the MGH Institutional Review Board. Human neutrophils were maintained in R10 (RPMI with 10% fetal bovine serum, L-glutamine, penicillin/streptomycin and HEPES) and grown at 37°C, 5%CO2 for the duration of the assay. Human NK cells were rested overnight in R10 supplemented with IL-15 at 37°C, 5%CO2 and maintained in R10 for the duration of the assay.

### Antibody-dependent cellular phagocytosis

Antibody-dependent cellular phagocytosis was assessed using a flow cytometry-based phagocytic assay *(47)*. Briefly, 1.0 μm, yellow-green fluorescent (505/515) FluoSpheres™ NeutrAvidin (Thermo Fisher) were coated with biotinylated S or HA, incubated with serum samples diluted 1:100, and breastmilk diluted at 1:10. The ability of samples to drive uptake of antigen-coated beads by THP-1 cells after overnight incubation was assessed by flow cytometry using the iQue (Intellicyt). Phagocytic scores were calculated as follows: (% yellow-green+ cells × yellow-green MFI)/100. A pool of SARS-CoV2 infected patients was used as a positive control, and a pool of non-infected patient serum and 1xPBS alone was used as a negative control. Samples were run in duplicate. Data reported is an average of all data points collected.

### Antibody-dependent neutrophil phagocytosis

Antibody-dependent neutrophil phagocytosis was measured by a flow cytometry-based assay *(48)*. Briefly, serum samples were diluted 1:100, while breast milk was diluted at 1:10. Samples were then allowed to form immune complexes with with 1.0 μm yellow-green fluorescent (505/515 nm) FluoSpheres™ NeutrAvidin (Thermo Fisher) beads coated with biotinylated antigen S or HA antigens. Neutrophils isolated from whole blood, obtained as mentioned above, using ACK Lysing Buffer (ThermoFischer) at room temperature (1:10). After lysis, remaining cells were counted and resuspended at 2.5×105 cells/ml in complete RPMI media. Cells were then incubated with bead/Ab mixture for 1 hr at 37°C and then stained with PacificBlue-conjugated anti-CD66b (Biolegend) in PBS. Finally, cells were fixed in 4% PFA and phagocytosis of beads by CD66b+ positive cells was measured by flow cytometry using the iQue (Intellicyt). A pool of SARS-CoV2 infected patients was used as a positive control, and a pool of non-infected patient serum and 1xPBS alone was used as a negative control. Samples were run in duplicate. Data reported is an average of data points collected from two donors.

### Antibody-dependent NK cell degranulation

Antibody dependent NK cell degranulation as described previously *(49)*. Briefly, antigen coated to 96-well ELISA plates were coated with 2 g/ml S or HA protein and incubated at 37°C for 2hrs and blocked with 5% BSA at 4°C overnight. NK cells were isolated from whole blood from healthy donors (as mentioned above) by negative selection using RosetteSep (STEMCELL Technologies) then separated using a ficoll gradient. NK cells were rested overnight in media supplemented with IL-15. Serum samples were diluted 1:50 and breastmilk at 1:5. After blocking, samples were incubated with antigen coated plates for 2 hrs at 37°C to allow immune complexes to form. After the 2 hr immune complex incubation, NK cells were mixed with anti-CD107a–phycoerythrin (PE)–Cy5 (BD), brefeldin A (10 μg/ml) (Sigma), and GolgiStop (BD), and added to antigen/ ab- coated plates for 5 hours at 37°C. The cells were stained for surface markers using anti-CD3 PacBlue (BD), anti-CD16 APC-Cy5 (BD), and anti-CD56 PE-Cy7 (BD) and fixed with PermA (Life Tech). NK cells were then permeabilized with Perm B (Life Tech) and stained with the cells were then analyzed by flow cytometry on the iQue (Intellicyt). NK cells were gates as CD3−, CD16+, CD56+ cells and NK cell activity was determined as the percent of NK cells positive for CD107a and MIP-1b. ADNKA was only performed on V2 samples and cord blood.

### Luminex

Serum samples were run in a customized Luminex assay to quantify the relative concentration of antigen-specific antibody isotype and subclass profiles. Carboxylated magplex-microspheres (Luminex) were coupled to antigens SARS-CoV-2 RBD (Sino Biological), SARS-CoV-2 S (Lake Pharma), SARS-CoV-2 N (Aalto Bio Reagents). SARS-CoV-2 S1 (Sino Biological), SARS-CoV-2 S2 (Sino Biological), pertussis pertactin (List Reagents) and a mix of HA A/Michigan/45/2015 (H1N1), HA A/Singapore/ INFIMH-16-0019/2016 (H3N2), B/Phuket/3073/2013 (Immunetech), using covalent NHS-ester linkages via EDC and NHS (Thermo Scientific) as described previously (Brown et. al 2017). To form immune complexes, appropriately diluted serum (1:100 for IgG2/3, 1:500 for IgG1, and 1:1000 for all other FcRs) and breastmilk (1:5 for all detectors), was added to the antigen-coupled microspheres, and plates were incubated overnight at 4°C, shaking at 700 rpm. The following day, plates were washed with 0.1% BSA 0.02% Tween-20. PE-coupled mouse anti-human detection antibodies (Southern Biotech) were used to detect antigen-specific antibody binding. For the detection of FcR binding, Avi-Tagged FcRs (Duke Human Vaccine Institute) were biotinylated using BirA500 kit (Avidity) per manufacturer’s instructions. Biotinylated FcRs were tagged with PE and added to immune complexes. Fluorescence was acquired using an Intellicyt iQue, and relative antigen-specific antibody titer and FcR binding is reported as Median Fluorescence Intensity (MFI).

### Univariate statistical analysis

For univariate data analysis, statistics were run using GraphPad Prism version 8.0. For functional assays, univariate analysis was performed comparing the three groups of women within each timepoint. Significance was determined by an ordinary one-way ANOVA for each antigen, with post-hoc Šídák’s multiple comparisons test for the reported adjusted p values. For Luminex assays, A p-value of less than 0.05 was considered significant. For comparisons between mother and either cord or breastmilk, Wilcoxon-matched pairs signed rank test was performed. To determine transfer ratios the cord serum or breastmilk value was divided by the respective mother’s value for each assay.

### Multivariate analysis

Multivariate analyses were performed in R (version 4.0.0) and Python (version 3.9.1). The data was then centered and scaled. For principal component analysis (PCA), an unsupervised dimensionality reduction technique, data was log10-transformed prior to centering and scaling.

For classification models, LASSO feature selection was performed using the “select_lasso” function in systemseRology R package (v1.0) (https://github.com/LoosC/systemsseRology). LASSO selection was performed 100 times, and only features that were chosen in 80% of the repetitions were used to build the downstream models. Partial least squares discriminant analysis (PLSDA) was performed using the LASSO-selected features. Multilevel partial least squares discriminant analysis (mPLSDA) *(50)* was performed for classification of maternal and cord samples. Model performance was evaluated by five-fold cross-validation. To evaluate model robustness, control models with permuted classifier labels or with random features were built 50 times. These control models were then cross-validated 10 times to determine model accuracy. P values were calculated as the probability that the true value was within the control distributions. For correlation networks, antibody features that had Bonferroni-adjusted p-values < 0.05 and a spearman |r|>0.75 were visualized within the networks.

## Supplementary Materials

Supplemental Figure 1. Similar antibody titers are induced in pregnant and lactating women following boost.

Supplemental Figure 2. Third-term vaccination results in an inefficient transfer of antibodies to the neonate, dependent on the time since vaccination to delivery.

Supplemental Figure 3. Boosting results in robust transfer of antibodies in breastmilk.

Supplemental Figure 4. Moderna and Pfizer vaccine-induced antibody profiles are similar at V1 but diverge at V2.

## Acknowledgements

We thank Bruce Walker, Nancy Zimmerman, Mark and Lisa Schwartz, and Terry and Susan Ragon for their support. We thank all members of the MGH Obstetric-Pediatric COVID-19 Biorepository Processing Team for assistance with specimen transport and processing. We thank the Brigham and Women’s/Massachusetts General Hospital Ob/Gyn residents and attendings for their support of the study. We acknowledge support from the Ragon Institute of MGH, MIT and Harvard and the Massachusetts Consortium on Pathogen Readiness (MassCPR). We thank SAMANA Kay MGH Research Scholars award for their support. Most importantly, we thank the participants for being part of the study.

## Funding

This work was supported by NIH-NHLBI grants K08HL1469630-02 and 3K08HL146963-02S1 (to K.J.G), NICHD grants 1R01HD100022-01, 3R01HD100022-02S2 (to A.G.E), the Ragon Institute of MGH, MIT, and Harvard and the MGH ECOR Scholars award, Nancy Zimmerman, an anonymous donor, and the Massachusetts Consortium on Pathogen Readiness (MassCPR), the NIH (3R37AI080289-11S1, R01AI146785, U19AI42790-01, U19AI135995-02, U19AI42790-01, 1U01CA260476 – 01, CIVIC5N93019C00052), the Gates Foundation Global Health Vaccine Accelerator Platform funding (OPP1146996 and INV-001650), and the Musk Foundation.

## Author Contributions

Conceptualization: G.A., A.G.E., C.A., M.A.E., E.A.D., and K.J.G

Investigation: C.A. and E.A.D.

Computational Analysis: C.A., C.D., and D.A.L.

Sample collection and processing: B.A., A.G.E., L.L.S., K.J.G., and R.M.D.

Supervision and Project Administration: E.A.B., M.A.E., A.F., and L.Y.

Writing – original draft: C.A., E.A.D., K.J.G., A.G.E., and G.A.

Writing – review & editing: All authors

## Competing Interests

Dr. Gray has consulted for Illumina, BillionToOne, and Aetion outside the scope of the submitted work. Dr. Alter is the founder of Seromyx Inc. Dr. Fasano reports serving as a cofounder of and owning stock in Alba Therapeutics and serving on scientific advisory boards for NextCure and Viome outside the submitted work. Dr. Elovitz reports serving as medical advisor for Mirvie. All other authors report no conflicts of interest.

## Data and materials availability

The dataset generated during this study is available upon reasonable request.

## Notes

### Competing Interest Statement

The authors have declared no competing interest.

