## Supplemental for "COVID-19 mRNA vaccines drive differential Fc-functional profiles in pregnant, lactating, and non-pregnant women"

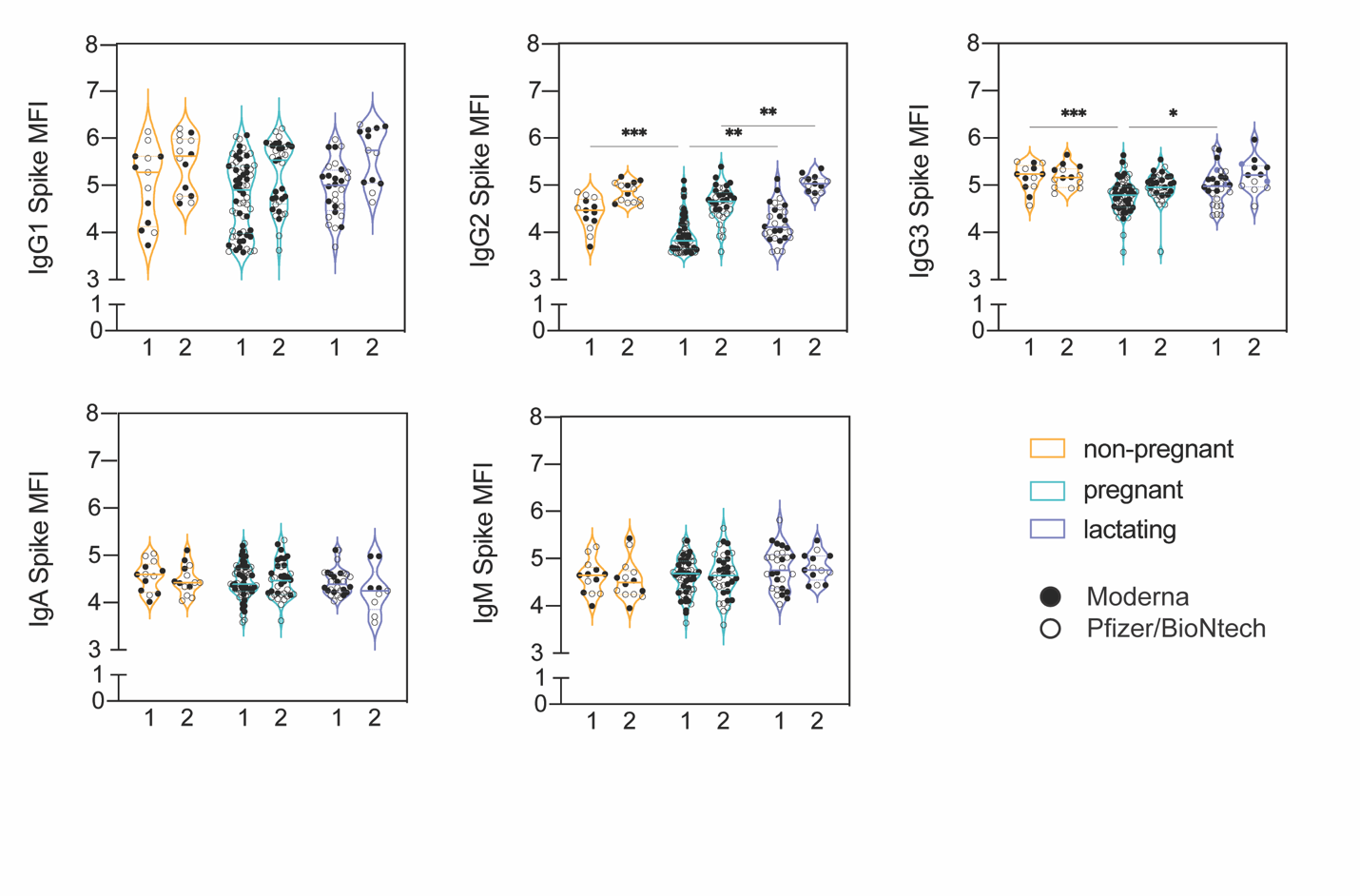

**Supplemental Figure 1. Similar antibody titers are induced in pregnant and lactating women following boost.**

The violin plots show the antibody titers for non-pregnant, pregnant, and lactating women at V1 (1) and V2 (2). The filled dots show the titer for women who received the Moderna vaccine, and outlines show the titer for women who received the Pfizer/BioNtech vaccine. Significance was only calculated between groups in the same time point and was determined by a one-way ANOVA with followed by posthoc Šidák’s multiple comparison test. * p <0.05, ** p < 0.01,*** p < 0.001, **** p < 0.0001.

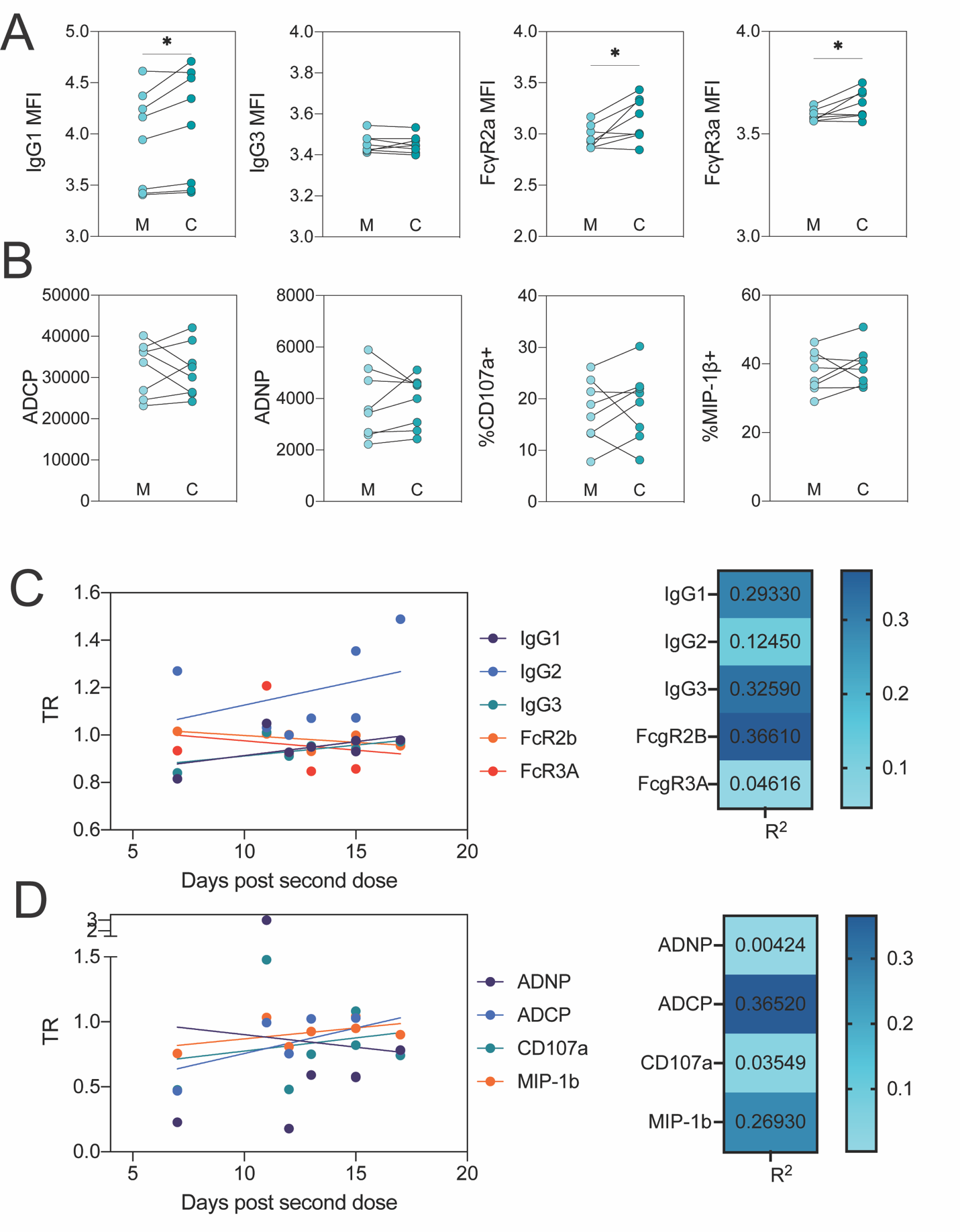

**Supplemental Figure 2. Third-term vaccination results in an inefficient transfer of antibodies to the neonate, dependent on the time since vaccination to delivery.**

**A.** The dot plots show the IgG1, IgG3, FcγR2a-binding and FcγR3a-binding titer against HA for maternal (M) and cord (C) blood. Lines connect maternal:cord dyads.

**B.** The dot plots show the ADCP, ADNP and ADNKA (CD107a and MIP-1β) functional titer against HA for maternal (M) and cord (C) blood. Lines connect maternal:cord dyads. Significance was determined by Wilcoxon-matched pairs signed rank test. * p <0.05, ** p < 0.01,*** p < 0.001, **** p < 0.0001.

**C.** Line graphs (right) show individual cord:maternal transfer ratios of spike specific antibodies at the time of delivery for IgG1, IgG2, IgG3, FcγR2B, and FcγR3A plotted in relation to the time post second dose. Linear regression R-squared values (left) for each feature plotted in on the right

**D.** Line graphs (right) show individual transfer ratios from mother to cord of spike specific antibodies at the time of delivery for IgG1, IgG2, IgG3, FcγR2B, and FcγR3A plotted in relation to the time post second dose. Linear regression R-squared values (left) for each feature plotted in on the right.

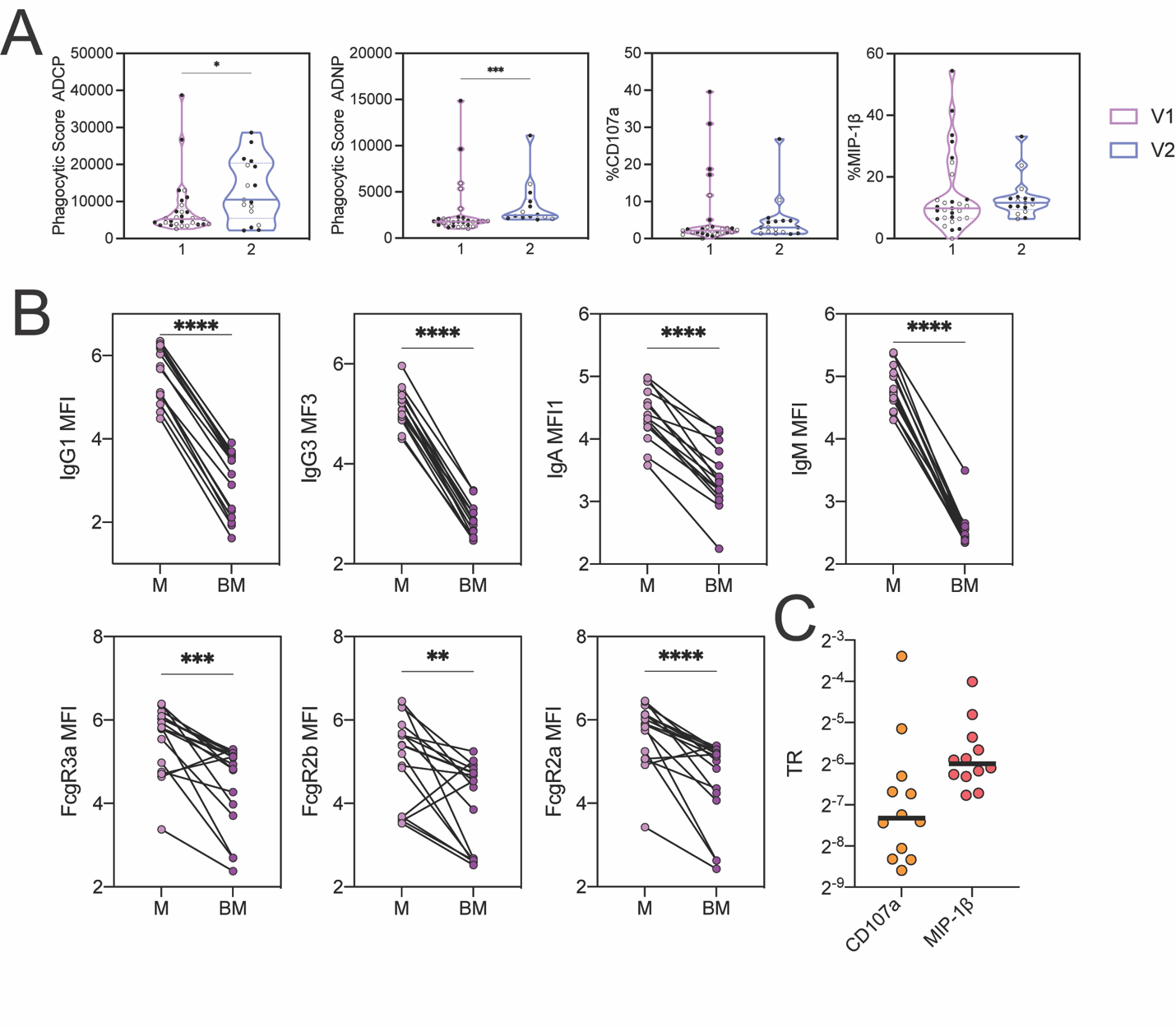
**Supplemental Figure 3. Boosting results in robust transfer of antibodies in breastmilk.**

**A.** The violin plots show the levels of ADCP, ADNP, and ADNKA (CD107a and MIP-1β) activity in breastmilk post-prime (V1, pink) and post-boost (V2, blue).

**B.** The dot plots show the IgG1, IgG3, IgA, IgM, FcγR2a-binding, FcγR2b-binding and FcγR3a-binding titers against SARS-CoV-2 spike at V2 in maternal serum (M) and breastmilk (BM). Lines connect matched maternal serum:breastmilk dyads. Significance was determined by Wilcoxon-matched pairs signed rank test. * p <0.05, ** p < 0.01,*** p < 0.001, **** p < 0.0001.

**C.** The dot plots show the transfer ratio (TR) (breastmilk/serum) of ADNKA (CD107a and MIP-1β) activity at V2.

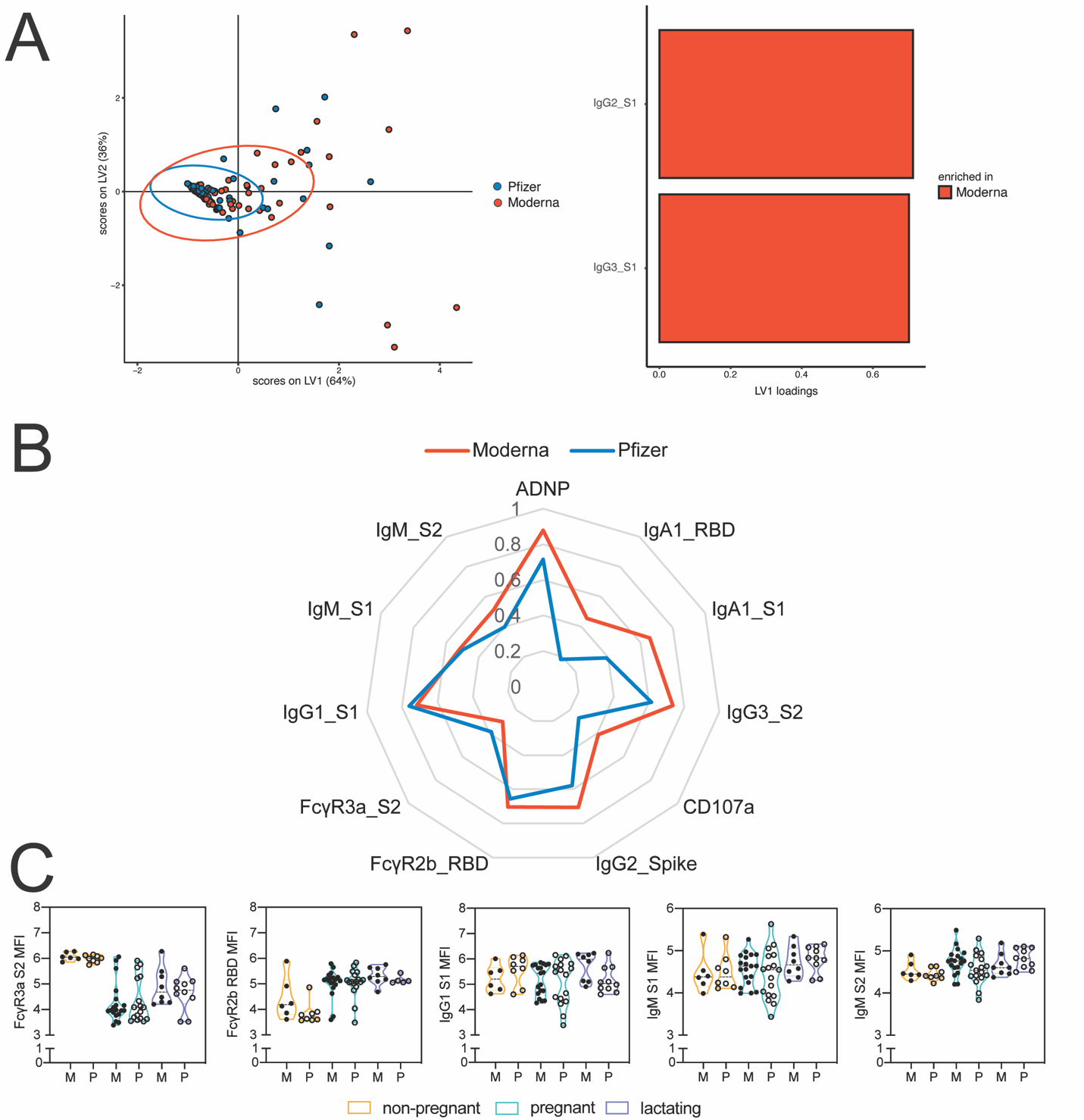

**Supplemental Figure 4**. **Moderna and Pfizer vaccine-induced antibody profiles are similar at V1 but diverge at V2.**

**A.** A LASSO PLSDA model was built on V1 data from all groups. The dot plot (top) shows the scores for each sample, with each dot representing a sample. The ellipses represent the 95% confidence interval for each group. The bar plot (bottom) shows the loadings of each LASSO-selected feature, where the color marks the group enrichment.

**B.** Spider plots show scaled values of each given antibody feature in each vaccine group.

**C.** The violin plots show differences in the top LASSO-selected features from Figure 4A in non-pregnant, pregnant and lactating women given either the Moderna (M, filled dots) or Pfizer/ BioNtech (P, outline dots). Significance was only calculated between groups in the same time point and was determined by a one-way ANOVA with followed by posthoc Šidák’s multiple comparison test. * p <0.05, ** p < 0.01,*** p < 0.001, **** p < 0.0001.
